# *De novo* evolution of antibiotic resistance to Oct-TriA_1_

**DOI:** 10.1101/2024.12.17.628969

**Authors:** Farhan R. Chowdhury, Laura Domínguez Mercado, Katya Kharitonov, Brandon L. Findlay

## Abstract

The rise of antimicrobial resistance as a global health concern has led to a strong interest in compounds able to inhibit the growth of bacteria without detectable levels of resistance evolution. A number of these compounds have been reported in recent years, including the tridecaptins, a small family of lipopeptides typified by the synthetic analogue octyl-tridecaptin A_1_. Hypothesizing that prior reports of negligible resistance evolution have been due in part to limitations in the laboratory evolution systems used, we have attempted to select for resistant mutants using a soft agar gradient evolution (SAGE) system developed by our lab. Following optimization of the media conditions by incorporation of the anti-synaeresis agent xanthan gum into the agar matrix, we successfully evolved high-level resistance to both octyl-tridecaptin A_1_ as well as the challenging lipopeptide antibiotic polymyxin B. Decreased tridecaptin susceptibility was linked to mutations in outer membrane proteins *ompC*, *lptD* and *mlaA*, with the effect of these genes confirmed through a mix of allelic replacement and knockout studies. Overall, this work demonstrates the robust evolutionary potential of bacteria, even in the face of challenging antimicrobial agents.

## Introduction

Antimicrobial resistance (AMR) is a major global health concern that threatens access to basic medical interventions. It is estimated that AMR was directly responsible for 1.27 million global deaths and contributed to 4.95 million deaths in 2019 (Murray *et al*., 2022), and it is currently projected that, if left unchecked, AMR will be responsible for 10 million deaths annually by 2050 (Jacobs, 2019).

Unfortunately, resistance to many potential new antibiotics can be found in bacterial pathogens before the drugs’ commercial release, due in part to cross-resistance between similar drug molecules (Bonomo *et al*., 2024). Studies that describe new antibiotics now often include adaptive laboratory evolution (ALE) experiments to determine rates of resistance or to elucidate the mechanism of action, with some antibiotics showing little to no resistance evolution (Mcguire *et al*., 1955; Ge *et al*., 1999; Ling *et al*., 2015; Cochrane *et al*., 2016; Stokes *et al*., 2020; Shukla *et al*., 2022). These latter antibiotics have attracted significant interest as promising candidates for next-generation antibiotic therapy, and may represent desirable “evolution-proof” or “resistance-proof” agents (Bell and MacLean, 2018; Upadhayay *et al*., 2023). However, the evolutionary resilience of many of these compounds has only been assessed through a limited array of ALE experiments, and has generally not been independently verified.

Tridecaptin A_1_ is one antibiotic against which laboratory evolution experiments have failed to describe *de novo* resistance. Originally isolated from *Paenibacillus* spp. (Shoji *et al*., 1978; Lohans *et al*., 2012), the tridecaptins are a group of non-ribosomal lipopeptides that act by selectively binding to the cell wall synthesis precursor lipid II of Gram-negative bacteria and dissipating the proton motive force (Cochrane *et al*., 2016). Their linear structure is readily accessible to solid phase peptide synthesis, allowing facile construction of tridecaptin analogues (Cochrane *et al*., 2014b; Ballantine *et al*., 2019). Best studied of these is octyl-tridecaptin A_1_ (Oct-TriA_1_), in which the chiral lipid tail is replaced with a low-cost octyl equivalent with no significant change in antimicrobial activity (Cochrane *et al*., 2014a). Tridecaptins are selective for Gram-negative bacteria, and their potent activity against the majority of the WHO’s priority pathogens list (WHO publishes list of bacteria for which new antibiotics are urgently needed, n.d.) makes them exciting antibiotic candidates. They have also been reported to be evolutionarily resilient, with Cochrane *et al*. finding no appreciable resistance to Oct-TriA_1_ following a 30-day laboratory evolution experiment (Cochrane *et al*., 2016).

We previously reported the ability of the soft agar gradient evolution (SAGE) system to rapidly generate resistance against antibiotics, including ones difficult to evolve in other platforms (Ghaddar *et al*., 2018). SAGE uses antibiotic gradients and bacteria’s natural propensity to swim through soft agar to select for antibiotic-resistant mutants. Unfortunately, the efficacy of SAGE is limited by synaeresis, the tendency of agar hydrogels to spontaneously shrink over time via continuous expulsion of solvent (Divoux *et al*., 2015). In SAGE this hinders bacterial motility (Croze *et al*., 2011) and limits experiments to ten days or less. We report here a new SAGE medium that is resistant to synaeresis. Supplemented with xanthan gum, a polysaccharide with excellent water binding capacity (Sánchez *et al*., 1995), this media has a reduced agar content and is suitable for month-long evolution experiments. We start by showing that resistance to the lipopeptide polymyxin B (PolB), an antibiotic that has proven difficult to evolve resistance to in SAGE (Ghaddar *et al*., 2018) and in other platforms (Trimble *et al*., 2016), can now be quickly achieved via SAGE. We subsequently use this medium to successfully generate resistance against Oct-TriA_1_ in *Escherichia coli* through a 27-day, maintenance free, SAGE experiment. Whole genome sequencing of evolved strains reveals mutations in phospholipid transport, outer membrane (OM) assembly and liopolysaccharide (LPS) biosynthesis. Notably, mutations in the *lptD* gene appeared consistently across resistant strains, implying its importance in resistance to Oct-TriA_1_. We then conduct further investigations into the role of *lptD*, *mlaA* and *ompC* mutations through allelic replacement studies, demonstrating their effect on Oct-TriA_1_ and other antibiotics minimum inhibitory concentrations (MICs), as well as their fitness costs.

## Results

### Standard SAGE medium fails to generate resistance to polymyxin B

To begin testing the ability of SAGE to generate resistance to lipopeptides, we attempted to evolve resistance to PolB in *Escherichia coli* K-12 substr. BW25113. However, we repeatedly failed to evolve resistance greater than 4-fold the initial value of 0.25 μg/mL. At low PolB concentrations ([PolB]_max_= 1.25 μg/mL, 5x MIC), cells quickly covered the plate and on isolation gave MICs that were 2x-4x that of the wildtype strain. However, the susceptibility of these mutants quickly reverted to wildtype (WT) levels upon subculturing in antibiotic-free media (data not shown), a feature consistent with the heteroresistance often observed with polymyxins like PolB and colistin (Hjort *et al*., 2016; Andersson *et al*., 2019; Liao *et al*., 2020). Growth in plates with a higher [PolB]_max_ (10 μg/mL, 40x MIC) failed to reach the end of the plates (Figure 1A, 1B). This behaviour is consistent with other ALE platforms, and PolB is known to be difficult to evolve resistance to via ALE (Trimble *et al*., 2016).

**Figure 1.**
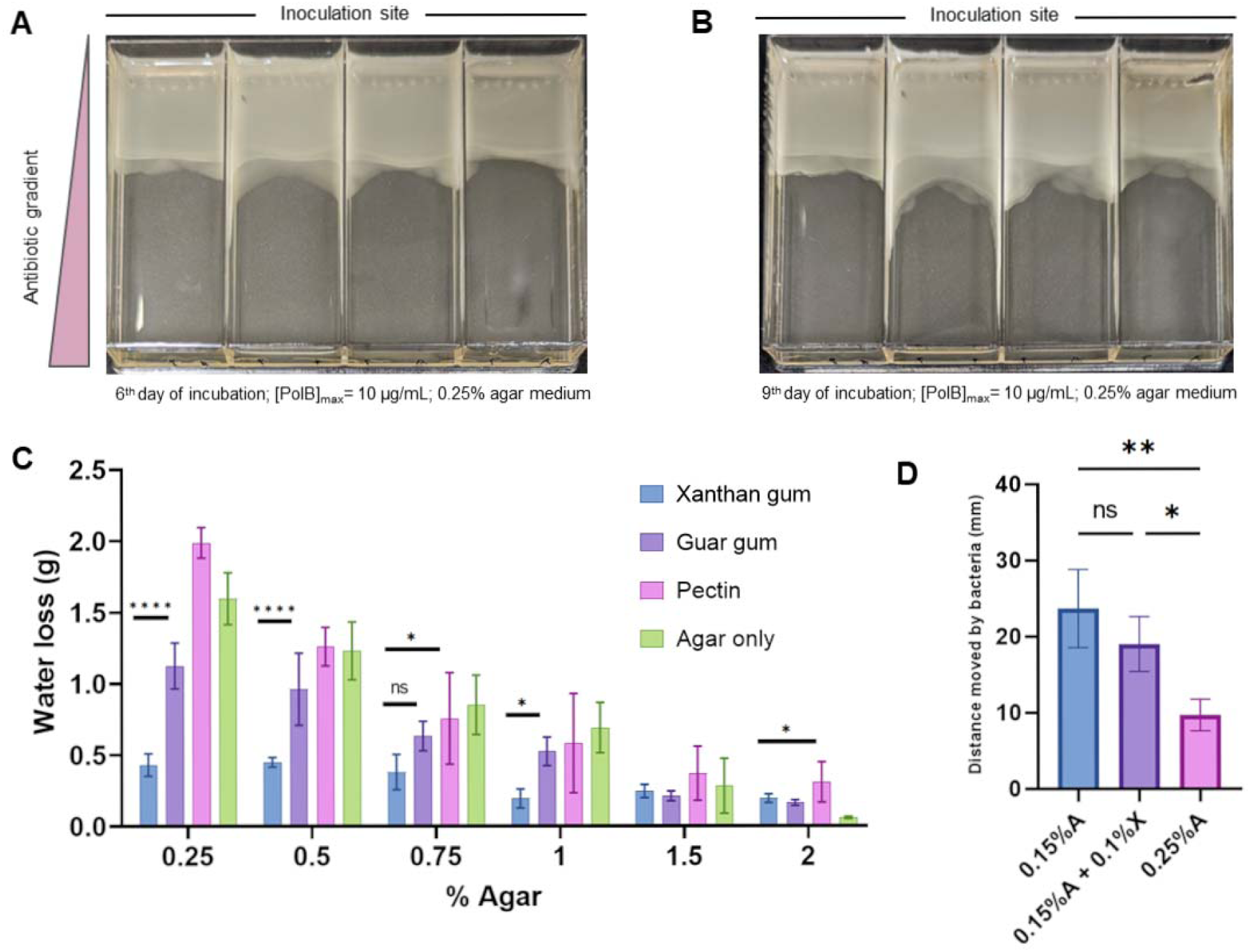
Synaeresis limits SAGE. (A) Cells in 0.25% agar-based PolB SAGE plates ([PolB]_max_= 10 μg/mL) remain stationary ∼30 mm from the inoculation site. (B) Further incubation results in only small movements of the bacterial front. (C) Xanthan gum outperforms all other additives tested for synaeresis-resistance across a range of agar strengths (n= 5). (D) Distance moved by bacteria in 0.15% agar medium (0.15%A), 0.15% agar + 0.1% xanthan gum medium (0.15%A + 0.1%X), and 0.25% agar medium (0.25%A). Bacteria traverse significantly higher distances in the 0.15%A + 0.1%X medium compared to the 0.25%A (n= 3). *p < 0.05, **p < 0,01, *** p < 0.001, ****p < 0.0001, one-way ANOVA with Fisher’s LSD test. Error bars represent SD.

SAGE evolutions rely on the ability of bacteria to move through the antibiotic gradients set up in soft-agar (0.25% agar) (Ghaddar *et al*., 2018). During incubation, synaeresis increases the effective agar concentration, reducing bacterial motility. We noticed that the bacterial front in PolB SAGE plates incubated for more than a week scarcely moved (Figure 1A, 1B), and hypothesized that synaeresis may be hindering the emergence of chromosomal mutations that confer stable PolB resistance. In line with this, a previous study reported that resistance to PolB in *E. coli* did not evolve for ∼6 days (in a liquid evolution platform), after which a rapid increase in resistance was seen (Yoshida *et al*., 2017). The authors proposed a two-step trajectory of resistance where heteroresistant bacterial populations leverage non-genetic mechanisms to withstand PolB stress at low concentrations, accessing stable chromosomal mutations only when the antibiotic concentrations increased (Yoshida *et al*., 2017). We thus set out to develop a SAGE medium more suitable for prolonged experiments.

### Xanthan gum supplementation reduces synaeresis in agar hydrogels

Polysaccharides like pectin, guar gum, and xanthan gum are able to form hydrogen bonds with water molecules, and are widely used as thickening agents in the food industry (Sánchez *et al*., 1995; Einhorn-Stoll, 2018). We hypothesized that the addition of these water-binding agents to agar gels may help slow down the synaeresis-driven remodeling of the agar matrix by resisting expulsion of water. We first confirmed that *E. coli* cannot utilize these polysaccharides as a carbon source (Supplementary Figure 1), then evaluated their effect on the synaeretic properties of agar gels. Each agent was separately added at 0.25% to agar strengths ranging from 0.25% to 2% (all percentages are in w/v), and the extent of synaeresis was evaluated via a modification of the method described by Banerjee *et al*. (Banerjee and Bhattacharya, 2011). Gels supplemented with xanthan gum achieved the highest reduction in water loss at all agar strengths tested (Figure 1C). While not a gelling agent itself, xanthan gum could replace a proportion of the agar while maintaining gel cohesion and limiting synaeresis (Supplementary Figure 2), though the medium became viscous at higher xanthan gum strengths.

Next, we tested the effect of addition of xanthan gum on bacterial motility in the SAGE medium. Addition of 0.1% xanthan gum to 0.15% agar had no statistically-significant effect on bacterial motility when compared to 0.15% agar alone, and both offered significant improvements in motility compared to the 0.25% agar medium. (Figure 1D, Supplementary Figure 3).

### Supplementation with xanthan gum enhanced the evolution of polymyxin B resistance

Moving forward, we opted to use a mixture of 0.2% xanthan gum and 0.15% agar (referred to from here on as XAM), a ratio which provided a balance of low viscosity in liquid state and high stability in the gel state, in place of the conventional 0.25% agar base used in SAGE. We found no difference between diffusion rates of malachite green in 0.25% agar and XAM (Supplementary Figure 3B), indicating that diffusion rates of antibiotics in XAM should be similar to that in the conventional medium.

To test the performance of the medium in SAGE, we set up a PolB SAGE plate with XAM ([PolB]_max_ in SAGE = 10 μg/mL, 40x MIC). We were able to generate stable PolB resistant mutants within 4 days in 2/4 SAGE lanes (MIC: 16 μg/mL, Figure 2A). We suspect that the increase in bacterial movement speed in xanthan gum-supplemented media (Figure 1D) reduced the time required for evolution in XAM-SAGE plates by allowing bacteria to reach PolB concentration that selects for stable genomic mutations earlier. By the time cells reached this concentration in the conventional medium, the medium may have already been too dry to allow movement.

**Figure 2.**
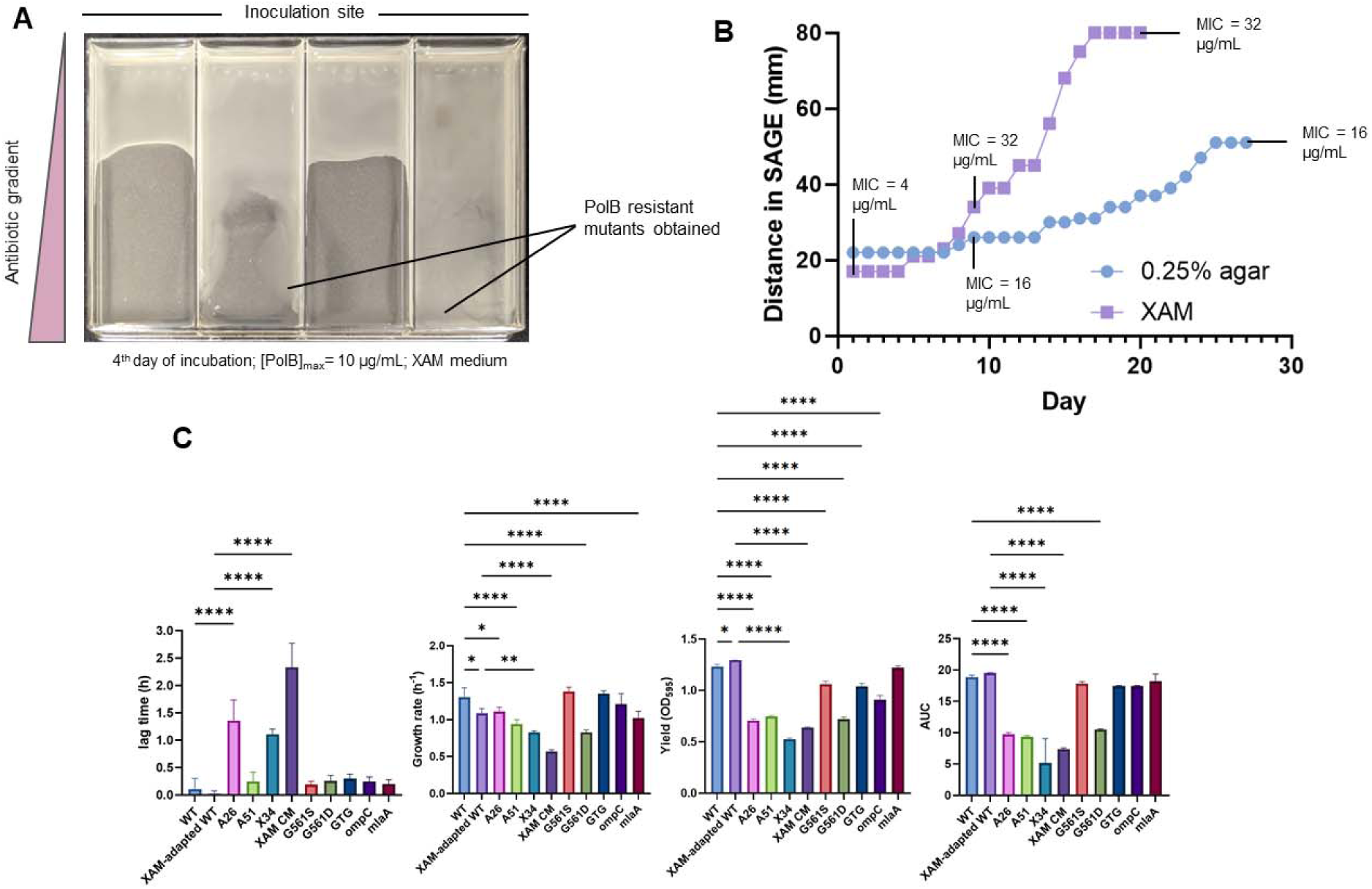
Evolution of antibiotic resistance. (A) Resistance to PolB emerged in 2 out of the 4 replicates in SAGE with XAM ([PolB]_max_= 10 μg/mL). (B) Distance moved by bacteria swimming through agar or xanthan gum/agar SAGE plates loaded with Oct-TriA_1_ at a max concentration of 40 μg/mL. Bacteria moved farther and faster in XAM. MIC of samples from several time points are labelled, with bacteria in XAM achieving a higher MIC (full list of MICs in Table 1). (C) Oct-TriA_1_ mutants are fitness impaired (n= 3). *p<0.05, **p<0.01, ***p<0.001, p<0.0001, one-way ANOVA with Bonferroni correction. For statistical comparisons, WT values were compared with XAM-adapted WT, A26, A51, G561S, G561D, GTG, ompC and mlaA, and XAM-adapted WT values were compared with XAM34 and XAM CM. Among these comparisons, only statistically significant differences are indicated by asterisks. Error bars represent SD. Results obtained from 3-9 replicates.

**Table 1:** Details of Oct-TriA_1_ mutants sampled from SAGE.

| Strain ID | Media in SAGE lane | Sampling day | Oct-TriA<sub>1</sub> MIC<br>($\mu\text{g/mL}$) |
| --- | --- | --- | --- |
| WT | - | - | 4 |
| XAM-adapted WT | XAM | - | 4 |
| A26 | 0.25% agar | 9 | 16 |
| A30 | 0.25% agar | 14 | 16 |
| A37 | 0.25% agar | 21 | 16 |
| A51 | 0.25% agar | 27 | 16 |
| XAM34 | XAM | 9 | 32 |
| XAM45 | XAM | 12 | 32 |
| XAM56 | XAM | 14 | 32 |
| XAM CM | XAM | 20 | 32 |

### Evolution of resistance to Oct-TriA_1_

We next sought to evolve resistance to Oct-TriA_1_ in XAM. We set up two SAGE lanes in parallel ([Oct-TriA_1_]_max_ = 40 μg/mL, 10x MIC), one with the conventional 0.25% agar medium, and the other with XAM. We followed the evolution of resistance by measuring the maximum distance moved by the bacterial fronts every 24 h (Figure 2B). Bacteria moved slowly through the 0.25% agar medium, traversing only about 60% of the lane (∼50 mm) by the end of 25^th^ day and remaining stationary for 3 additional days before the experiment was stopped (Figure 2B). The small distances moved and thinning agar gel made sampling from this medium beyond 7 days challenging. In contrast, bacteria in the XAM lane moved large distances after breaking free from the initial inhibitory Oct-TriA_1_ concentration (Day 7), covering the entire lane by the end of day 17 (Figure 2B). The XAM gel also appeared to have retained significantly more water than the agar-based gel at the end of the experiment (data not shown). Samples were collected whenever significant bacterial movement was detected (Methods). We tested the MIC of samples A26, A30, A37, A51, XAM34, XAM45, XAM56 and XAM CM (‘A’ and ‘XAM’ in the sample IDs denote samplings from 0.25% agar lane and XAM respectively, and the numbers denote the distance in millimeters from the inoculation zone to where cells were sampled; CM = cells extracted from the end of lane, ∼75 mm) (Table 1). The MIC of A26 and XAM34, both from the 9^th^ day of incubation, showed that resistance emerged early, and appeared to remain constant throughout the rest of the experiment (Table 1). Since standard MIC assays are based on 2-fold dilution steps, we suspect that small increases in MIC might have occurred after the initial increase but could not be resolved via the MIC assays. Overall, mutants from the 0.25% agar-based medium exhibited up to 4x increase in MIC, compared to an 8x increase in XAM (Table 1).

Next, we compared the fitness of the early and endpoint Oct-TriA_1_ mutants to that of the WT parent strain (Figure 2C). The WT *E. coli* used for generating mutants from the 0.25% agar lane was pre-adapted to this SAGE medium as previously described (Ghaddar *et al*., 2018). To account for any changes in fitness due to adaptation to XAM, we also passaged the WT strain through antibiotic-free XAM 3 times to produce a XAM-adapted WT strain (Materials and Methods). All evolved mutants showed longer lag times (though differences with A51 did not reach statistical significance) and lower growth rates, yields and AUCs (area under the growth curves), indicating that Oct-TriA_1_ resistance imposed a large fitness cost (Figure 2C). In general, the XAM-generated mutants showed larger fitness deficits, even though the fitness of the XAM-adapted WT was comparable to the WT in every metric measured. Interestingly, the XAM CM strain showed a clear diauxic growth pattern (Supplementary Figure 4). This strain harbored a deletion in the *nuo* operon, which codes for a NADH/ubiquinone oxidoreductase that shuttles electrons from NADH into the electron transport chain (Prüss *et al*., 1994; Van den Bergh *et al*., 2022). When cells grow in the presence of glucose, they excrete acetate (Shimada and Tanaka, 2016). As cells deplete glucose from media, they switch to uptaking acetate (Shimada *et al*., 2021), shifting from glycolysis to TCA cycle and gluconeogenesis (Shimada and Tanaka, 2016). In *nuo* mutants, high NADH/NAD^+^ ratios inhibit enzymes involved in the TCA cycle, drastically slowing growth and potentially giving rise to the diauxic growth pattern we observed (Prüss *et al*., 1994; Shimada *et al*., 2021). However, what causes subsequent resumption of growth during diauxie is unclear (Chu and Barnes, 2016; Salvy and Hatzimanikatis, 2021).

### Genetic analysis of Oct-TriA_1_-resistant mutants

XAM34 had an MIC eight times that of the wildtype *E. coli* BW25113 and had eight mutations: three non-synonymous, three intergenic, and two frameshift insertions (Table 1, Figure 3). XAM_CM, drawn from later in the same SAGE plate, had the same MIC and ten mutations: four non-synonymous, two intergenic, three frameshift insertions, and one frameshift deletion. Two of these were identical: a five-base deletion in *yddW* and a E26K mutation in *rpoD*. A26 had two nonsynonymous mutations and the same five-base deletion in *yddW,* while A51 had four non-synonymous mutations, one intergenic mutation, one insertion, and three frameshift mutations (including the five-base deletion in *yddW*).

**Figure 3.**
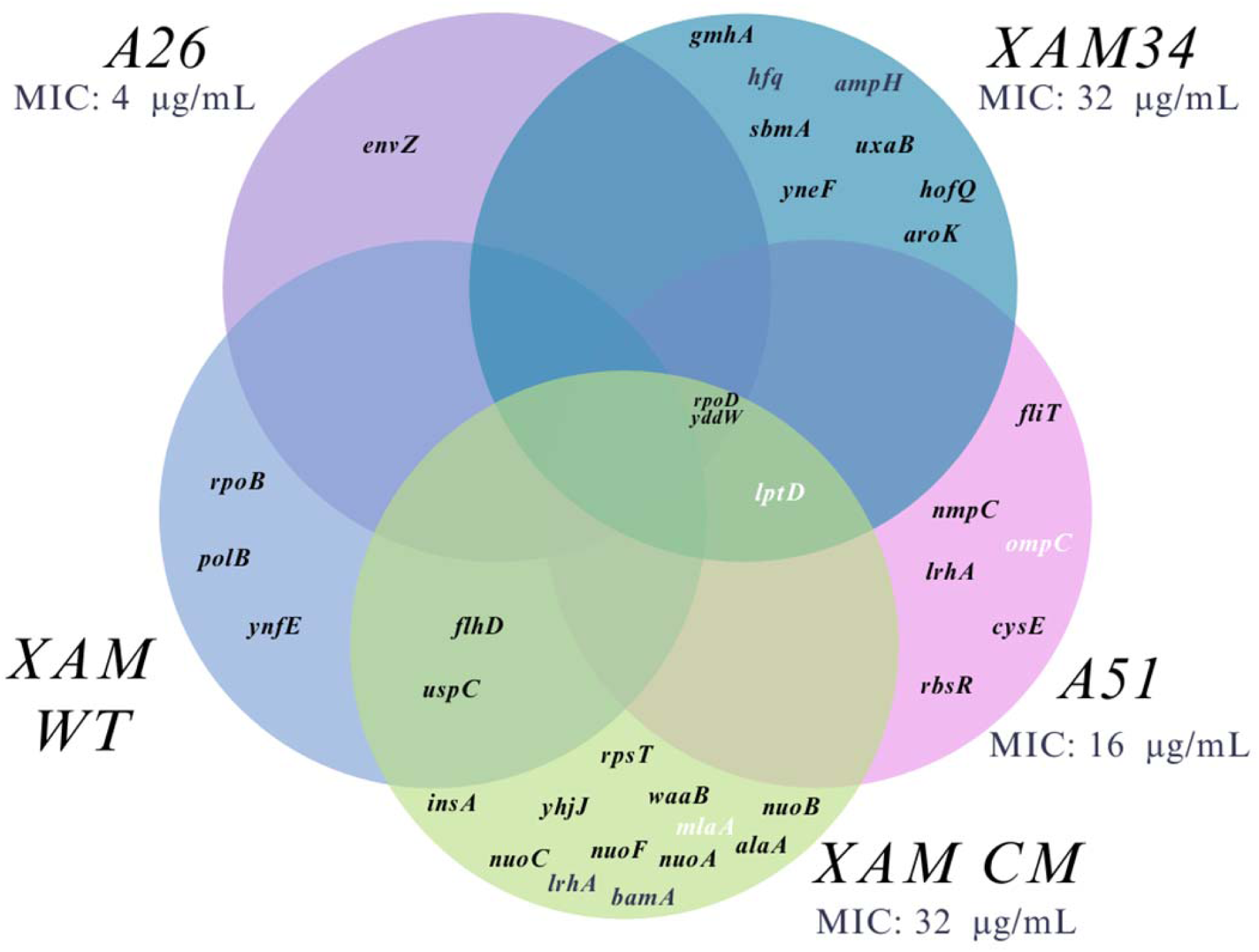
Mutations identified in the evolved strains. A26 and A51 represent mutations observed in cells isolated at 26 mm and 51 mm in agar media; XAM 34 and XAM CM cells isolated at 34 mm and 75 mm, respectively in xanthan gum media. The highlighted mutations were selected for allelic exchange.

To separate possible resistance mutations from those that might be associated with adaptation to the ALE conditions, we sequenced the XG-adapted WT for comparison with the XAM-generated Oct-TriA_1_ mutants. Only one mutation overlapped with the antibiotic-exposed samples; an intergenic mutation in *flhD* ← / → *uspC* that was also found in XAM CM. The XAM evolved strain carried a G→T mutation in 261/519 while the XAM CM carried a C→A in 263/+44. *flhD* is involved in flagellar type II transcription activation and *uspC* is a universal stress protein. Changes in *flhD* expression may alter swimming speed, enhancing movement through the soft agar plates (Barker *et al*., 2004; Wang and Wood, 2011; Lee and Park, 2013). Other mutations observed were in *rpoB*, *ynfE* and *polB* genes, genes not mutated in any of the antibiotic-exposed strains.

All strains with MICs higher than the wild type carried a single nucleotide polymorphism in *lptD,* creating LptD G561S (A51, XAM 34) or G561D (XAM CM). This was also the only gene to contain two mutations, with A51 having a further (GTG)3→2 deletion in nucleotides 705 707. LptD is an integral component of the Lpt complex, which is essential for the assembly and transport of LPS to the outer membrane of Gram-negative bacteria (Chng *et al*., 2010). Mutations were also observed in two other genes linked to LPS biosynthesis: X34 contained a SNP in *gmhA*, which encodes a phosphoheptose isomerase that produces the D-*glycero*-D-*manno*-heptose 7-phosphate found in the core of LPS (Taylor *et al*., 2008), while XAM CM contained a frameshift mutation in *waaB,* which encodes a galactosyltransferase that appends galactose to that core (Qian *et al*., 2014). Similar to polymyxin B (Trimble *et al*., 2016), tridecaptin A_1_ engages with LPS in the outer membrane to facilitate uptake into the intermembrane space and access its target (Cochrane *et al*., 2016), and these mutations strongly suggest that tridecaptin resistance is conferred by alterations in LPS structure.

Three other mutations had clear ties to the bacterial outer membrane: *mlaA* directly regulates outer membrane composition and was heavily truncated in XAM CM (MlaA W59*), while the porin gene *ompC* is implicated in both maintenance of outer membrane integrity and drug uptake (Chong *et al*., 2015; Choi and Lee, 2019). Notably, OmpC was altered both directly through a nonsynonymous *ompC* SNP (OmpC N47S) in A51 and indirectly through a nonsynonymous SNP in the *omp* regulator *envZ* (EnvZ R253S) in A26. A SNP was also observed in *bamA* (BamA L501Q; XAM CM), part of the BAM complex (Lehman and Grabowicz, 2019). BamA is responsible for inserting β-barrel proteins into the OM (Lehman and Grabowicz, 2019). Mutations in *bamA* have been related to resistance to drugs targetting this OM protein (Hart *et al*., 2019; Luther *et al*., 2019; Kaur *et al*., 2021). In the case of darobactin, resistance mutations in *bamA* also result in loss of virulence (Huang *et al*., 2019).

Gene ontology enrichment analysis mapped the remaining mutations to several key pathways, including those related to respiratory electron transport mechanisms (Supplementary Table 1). This suggests adaptive changes in electron transport and ATP synthesis, in addition to alterations in outer membrane assembly and biosynthesis.

### Allelic exchange in genes involved in phospholipid transport and outer membrane assembly confirmed their involvement in resistance to Oct-TriA_1_

To investigate the effect of mutations in genes associated with phospholipid transport, we introduced the observed mutations in *lptD* into *E. coli* BW25113 via allelic exchange. This was carried out using CRISPR-Cas9/λ-Red assisted recombineering as previously described (Reisch and Prather, 2015; Reisch and Prather, 2017). Concurrently, knockouts in *ompC* and *mlaA* were obtained from the Keio collection (Baba *et al*., 2006). The effect of these alterations was assessed via MIC assays, revealing that all five alterations increased the ancestral strain’s MIC against Oct-TriA_1_ two-fold (Supplementary Figure 5). The effect of the mutations on fitness was more variable. None of the mutations altered lag times during growth in MHB, but significant deviations were observed in both growth rates and max OD (Figure 2C). The effect of the LptD mutations on fitness varied by both site and type of mutation. Despite halving susceptibility towards OctTriA1, the extra GTG repeat had no effect on bacterial fitness, while the LptD G561D mutation was much more detrimental than the LptD G561S mutation. As no single mutation increased resistance or impaired fitness to the levels observed in XAM CM, a combination of costly mutations appears to be required for high-level resistance.

## Discussion

In this study we demonstrate the first *de novo* evolution of resistance to Oct-TriA_1_, with the effect of putative resistance-conferring mutations confirmed through allelic exchange. Further, we have improved the SAGE system through the incorporation of the thickening agent xanthan gum, extending the potential duration of experiments from a week to a month and enhancing selection rates. This modified system was also much more effective at selecting mutants resistant to polymyxin B, an antibiotic that is often difficult to target with other ALE systems.

In line with resistance to other D-amino acid-containing non-ribosomal peptides (Li *et al*., 2018), resistance in the native producers of tridecaptins is mediated via hydrolytic D-stereospecific peptidases (Bann *et al*., 2021). In contrast, the mutations we have observed are largely in genes coding for LPS biosynthesis and outer membrane homeostasis. These pathways are essential to bacterial growth, as well as interactions with the immune system, nutrient acquisition, and toxin susceptibility (Liu *et al*., 2012; Phan and Ferenci, 2017; Simpson and Trent, 2019). As a result, it is unsurprising that the resistant strains we generated had significantly impaired fitness (Figure 2C). Similar results have been observed with other membrane-interacting antibiotics, like the polymyxins. In many pathogens mutations that confer colistin resistance significantly impairs fitness and/or virulence (Wang *et al*., 2022), though acquisition of the plasmids encoding colistin resistance factor *mcb-1* has a much smaller impact (Tietgen *et al*., 2018).

The factors that underpin widespread, high-level resistance are not fully understood. When evaluating evolution potential there has been a strong tendency to focus on the rate by which resistance emerges, either through mutation rate studies or ALE (Martinez and Baquero, 2000; Cirz *et al*., 2005; Ling *et al*., 2015; Cochrane *et al*., 2016; Sommer *et al*., 2017; Stokes *et al*., 2020; Martin *et al*., 2020). This work suggests that the nature of the mutations should also be taken into account. Each of the mutations in *lptD, mlaA,* and *ompC* altered the octyl-tridecaptin A_1_ MIC two-fold, with little overlap between strains (Figure 3, 4). Given the overall change in susceptibility following SAGE was 32-fold, high-level resistance likely resulted from a combination of multiple mutations rather than from any single mutation. SAGE is well-suited to the serial acquisition of small-impact mutations (Ghaddar *et al*., 2018), potentially explaining why it was successful when attempts to evolve resistance via serial passage through liquid culture failed (Cochrane *et al*., 2016).

Xanthan gum was able to significantly reduce synaeresis and allow SAGE experiments to extend beyond their initial limit of 7-10 days, and this media may have utility outside ALE. Syneresis causes loss of growth-promoting properties of media when cultivating slow-growing bacteria and fungus (Laserna *et al*., 1981; Divoux *et al*., 2015; Savinova *et al*., 2023). Addition of a water-binding agent like xanthan gum may preserve these properties, allowing extension of those experiments as well.

Against the rising prevalence of antibiotic resistance, “evolution-proof” or “resistance-proof” are very appealing targets (Bell and MacLean, 2018; Upadhayay *et al*., 2023). Their discovery could greatly alleviate the growing AMR crisis, carving a path forward for the use of antibiotics for decades to come. However, since the discovery of sulfa drugs hundreds of antibiotics have entered clinical use, with pathogens evolving resistance to each and every one of them (Bell and MacLean, 2018). This work underscores the genetic flexibility of bacteria, and highlights the need for stringent evolution studies during the development and discovery of new antibiotics. If resistance is to emerge, we would do well to study it *in vitro* before its appearance in pathogens.

## Materials and Methods

### Bacterial Strain and Growth Conditions

*E. coli* K-12 substr. BW25113 and all subsequent resistant mutants were grown in cation-adjusted Mueller Hinton Broth (MHB 2) media at 37 °C. Liquid cultures were shaken at 250 RPM, while agar cultures were grown in a static incubator.

### Oct-TriA_1_ synthesis

Oct-TriA_1_ synthesis was performed as described by Cochrane *et al*. (Cochrane *et al*., 2014b), with the following modifications. Briefly, in a manual peptide synthesizer 120.5 mg of Wang resin pre-loaded with Fmoc-Alanine at a loading of 0.6 mmol/g was swelled in dimethylformamide (DMF). The protecting group was cleaved with two twenty-minute treatments of 4:1 DMF:4-methylpiperidine. The beads were then washed three times with DMF, once with dichloromethane (DCM), then one final time with DMF. The next residue in the series was then added in 3x excess, alongside HATU (3x excess) and diisopropylethylamine (DIPEA) (8x excess). Coupling was carried out for one hour, at which point the beads were washed as above and the Fmoc protecting group once more cleaved. This cycle was repeated for each of the peptides, with Fmoc-Glu(OtBu)-OH, Fmoc-D-Ser(OtBu)-OH, Fmoc-Dab(Boc)-OH, and Fmoc-D-Dab(Boc)-OH used for the residues with reactive side chains. The complete peptide was cleaved from the resin with a 95:2.5:2.5 solution of trifluoroacetic acid (TFA):deionized water:triisopropylsilane for 2 hours. The cleavage solvent was removed on a rotary evaporator, and the crude material was triturated three times with diethyl ether. The solid residue was then purified to homogeneity on an Agilent 1100 preparative HPLC system, using an XBridge BEH C18 OBD prep column (5 µm, 25 x 250 mm) and the following water/acetonitrile gradient system.

| Time | Acetonitrile (%) |
| --- | --- |
| 0 | 5 |
| 1 | 5 |
| 3 | 20 |
| 23 | 55 |
| 31 | 95 |
| 36 | 95 |

Peaks eluting around 13.2 min across multiple runs were pooled, and the identity of the peptide was confirmed via high resolution mass spectrometry on an Orbitrap LTQ Velos.

### SAGE evolutions

SAGE plates were prepared and inoculated as described previously (Ghaddar *et al*., 2018). For SAGE plates made with XAM, MH media + 0.15% agar was first stirred in a flask on a hot plate and stirrer on high for 5-10 minutes. 0.2% xanthan gum was then slowly added to the stirring liquid and the mixture was allowed to stir for 2-3 minutes before autoclaving. This medium was melted on demand prior to use in SAGE plates. As needed cells were extracted from SAGE plates by pipetting up 20 μL of the gel and transferring it into 5 mL MH media for culturing. Overnight growth was streaked on MH plates and single colonies were used to prepare glycerol stocks.

### MIC Assays

MICs were determined via broth microdilution, following CLSI guidelines (CLSI, 2018). Briefly, antibiotics were serially diluted in 96-well plates and mixed with bacteria at a final concentration of 5 x 10^5^ CFU/mL. Plates were incubated at 37 °C without shaking for 16-20 h, and the MIC was recorded as the lowest concentration that visibly inhibited growth.

### Synaeresis tests

Water loss from different gel mixtures was measured as described by Banerjee et al. (Banerjee and Bhattacharya, 2011) with the following modifications. Agar concentrations ranging from 0.2-2% were first stirred in a flask on a hot plate for 5-10 minutes. 0.25% xanthan gum, guar gum or pectin was then slowly added to the stirring liquid, which was allowed to stir for 2-3 minutes. Flasks were transferred to a 37 °C shaker and shaken overnight at 250 rpm to produce a smooth, homogenous mixture. The flasks were then autoclaved, and 20 ml of each liquid was transferred to 50 mL centrifuge tubes. Tubes were allowed to cool at room temperature, then stored at 4 °C overnight. Initial masses of the tubes were recorded (∼30 g on average) before centrifugation at 1000 rpm for 30 mins at 25 °C. Centrifugation broke the gel structure, making it difficult to decant water out of the tubes without losing gel mass. To extract the free liquid tubes were instead left upright with their caps open and a folded filter paper was used to wick away the water over 30 minutes. The filter papers were then carefully removed to minimize the loss of gel mass, and the final tube masses were recorded. Water loss was calculated as the difference between the initial and the final masses of the tubes.

### Bacterial Motility tests

Bacterial migration speeds on different gel compositions were measured as described by Croze et al. (Croze *et al*., 2011). The media were prepared as described above. 30 mL of each mixture was then poured in separate petri dishes, and the plates were left to set overnight at room temperature. 2 µL of overnight bacterial culture was placed on the center of each petri dish, and the inoculum was allowed to dry/absorb for an hour. 9 mL of mineral oil was overlaid on each plate, and all plates were then incubated at 37 °C without shaking, lid side up. The diameter of growth was measured 6 h post incubation.

### Generation of the XAM-adapted WT strain

12 mL of antibiotic-free XAM was poured in a SAGE lane and allowed to cool and solidify. 50 μL of overnight WT bacterial culture was then inoculated on one side and the inoculum was allowed to dry/absorb for 30 minutes. 2.5 mL of mineral oil was overlaid on the gel, and the plate was incubated at 37 °C. The next day, cells were extracted from the end of the lane as described above, then grown overnight. These cells were used to inoculate a second antibiotic-free XAM lane and the whole process was repeated. Following three consecutive passes, cells were streaked on agar, and a single colony was designated as the XAM-adapted WT strain.

### Fitness measurements

1 μL of overnight bacterial culture was added to 99 μL of MH broth in 96 well plates. Lids were treated with 0.05% Triton X-100 in 20% ethanol to reduce fogging (Brewster, 2003). Absorbance readings (595 nm) were recorded using a plate reader at 5 min intervals for 24 h (Tecan Sunrise). Area under the growth curves were calculated in GraphPad Prism. All other metrics were generated using Dashing Growth Curves (Reiter and Vorholt, 2024).

### WGS and variant calling

Whole genomes were extracted using a bacterial genomic DNA extraction kit following the manufacturer’s instructions (Bio Basic Inc, Cat: BS624). Whole genome sequencing was performed at SeqCenter using the Illumina NovaSeq X Plus sequencer, which generated 2x151 bp paired-end reads (Illumina Whole Genome Sequencing, n.d.). The Breseq v0.37.1 pipeline was used for variant calling with bowtie2 v2.4.5 and R v4.2.2 (Barrick *et al*., 2014; Illumina Whole Genome Sequencing, n.d.).

### Gene ontology enrichment analysis

Mutations observed in all of the evolved strains were analyzed for enrichment of gene ontology groups using the ShiniGO package v0.741 (Ge *et al*., 2020). The p-value cut-off for the False Discovery Rate (FDR) was set to 0.05 against *E. coli* MG1655, a K12 strain. Several previously reported differences between MG1655 and BW25113 were identified and excluded from the analysis (Grenier *et al*., 2014); most notably deletion of the *araBAD* and *rhaDAB* operons, replacement of a section of the *lacZ* gene with four *rrnB* terminators, and a frameshift mutation in *hsdR* that causes a premature stop codon.

### Allelic exchange mutant generation

Allelic exchange of the selected mutated genes was carried out using the no-SCAR (Scarless Cas9 assisted recombineering) method, as previously described (Reisch and Prather, 2015; Reisch and Prather, 2017). In short, retargeting of the pKDsgRNA plasmid was constructed for the *lptD* gene region of interest through CPEC cloning in a way that the mutation would disrupt the PAM site or the 12 bp seed region. Cas9 counterselection was achieved by sequentially transforming pCas9cr4 and the retargeted pKDsgRNA and electroporating dsDNA containing the desired mutation. Following induction of λ-Red and Cas9, the successful mutants were verified and the plasmids were cured of the plasmids to render them susceptible to Chloramphenicol and Spectinomycin.

### Keio collection strains Kan cassette curing

Keio collection strains were cured of the kanamycin resistance cassette through FLP-recombinase-mediated recombination, using the pCP20 plasmid as previously described (Datsenko and Wanner, 2000; Baba *et al*., 2006). Subsequent curing of temperature sensitive pCP20 plasmid yielded Kan^S^, Amp^S^ for MIC determination.

#### Strains

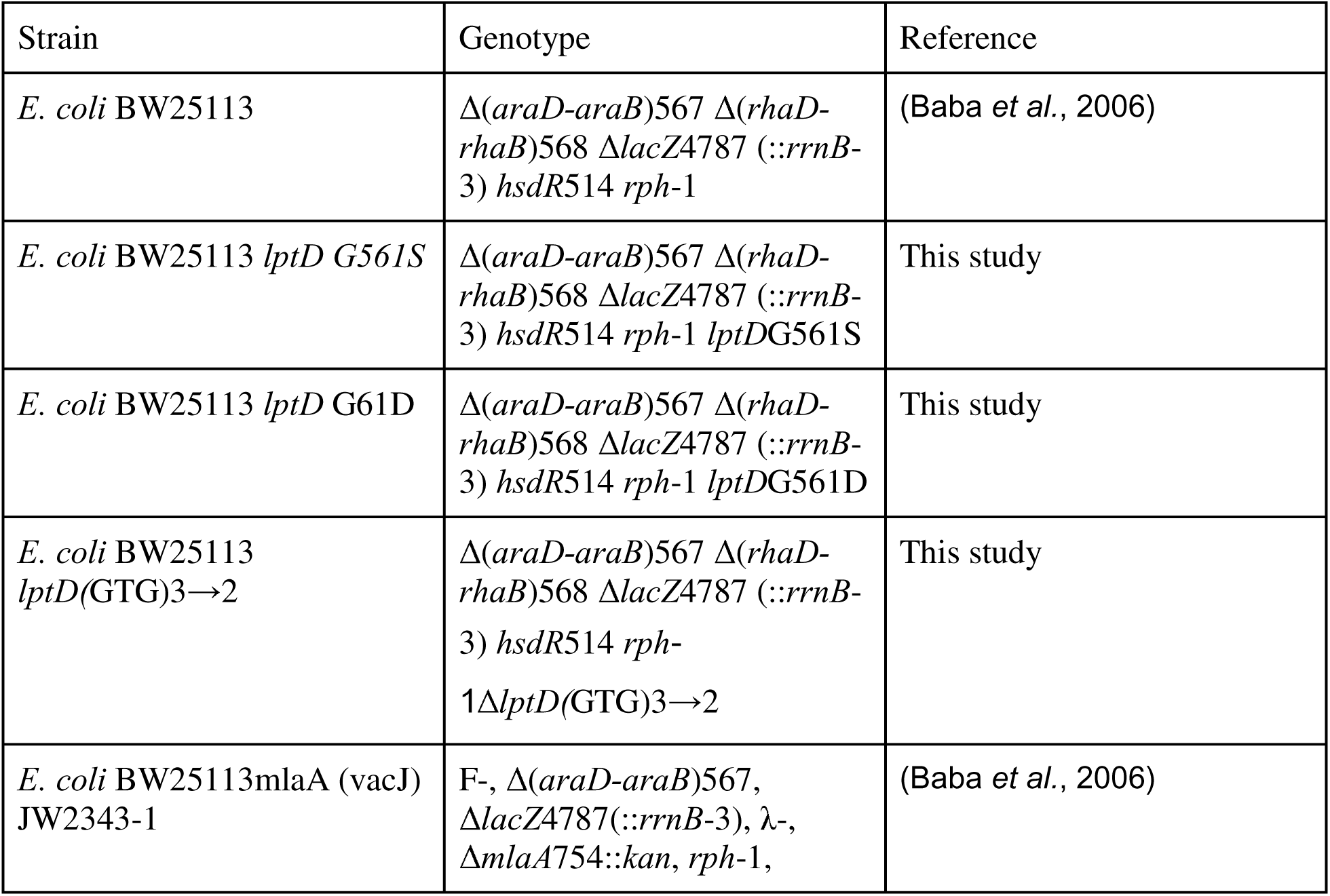
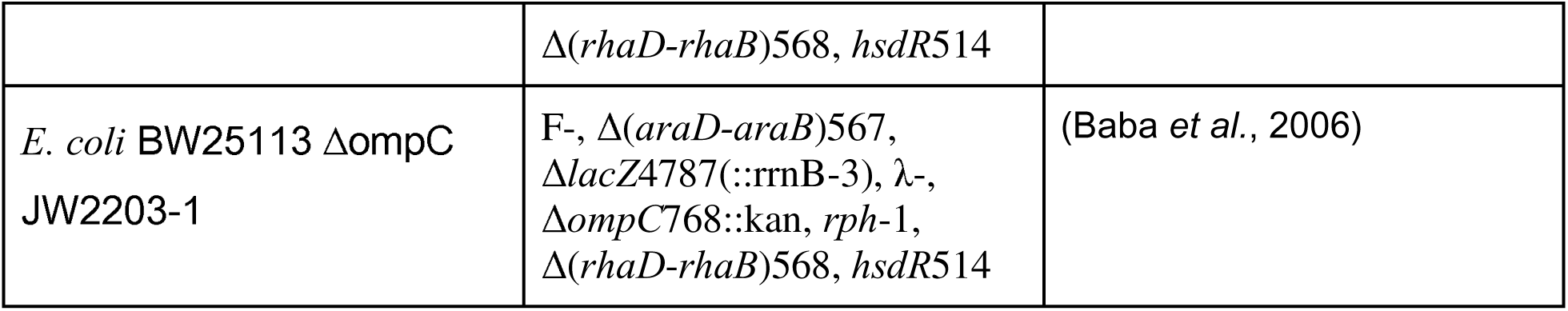

| Strain | Genotype | Reference |
| --- | --- | --- |
| *E. coli* BW25113 | $\Delta(araD-araB)567 \Delta(rhaD-rhaB)568 \Delta lacZ4787 (::rrnB-3) hsdR514 rph-1$ | (Baba *et al.*, 2006) |
| *E. coli* BW25113 *lptD* G561S | $\Delta(araD-araB)567 \Delta(rhaD-rhaB)568 \Delta lacZ4787 (::rrnB-3) hsdR514 rph-1 lptDG561S$ | This study |
| *E. coli* BW25113 *lptD* G61D | $\Delta(araD-araB)567 \Delta(rhaD-rhaB)568 \Delta lacZ4787 (::rrnB-3) hsdR514 rph-1 lptDG561D$ | This study |
| *E. coli* BW25113 *lptD*(GTG)3→2 | $\Delta(araD-araB)567 \Delta(rhaD-rhaB)568 \Delta lacZ4787 (::rrnB-3) hsdR514 rph-1 \Delta lptD(GTG)3\rightarrow 2$ | This study |
| *E. coli* BW25113 mlaA (vacJ) JW2343-1 | F-, $\Delta(araD-araB)567, \Delta lacZ4787 (::rrnB-3), \lambda$-, $\Delta mlaA754::kan, rph-1,$ | (Baba *et al.*, 2006) |
| | $\Delta(rhaD-rhaB)568, hsdR514$ | |
| *E. coli* BW25113 $\Delta ompC$<br>JW2203-1 | F-, $\Delta(araD-araB)567$,<br>$\Delta lacZ4787(::rrnB-3)$, λ-,<br>$\Delta ompC768::kan$, *rph-1*,<br>$\Delta(rhaD-rhaB)568, hsdR514$ | (Baba *et al.*, 2006) |

## Data Availability

WGS data is available under the NCBI sequence Read Archive BioProject: PRJNA1131392; BioSample accession numbers SRR29693652, SRR29693651, SRR29693650, SRR29693649, SRR29693648 and SRR29693647.

## Declaration of generative AI and AI-assisted technologies in the writing process

During the preparation of this work the authors used ChatGPT Plus (OpenAI, GPT-4) to generate alternative phrasings for some complex sentences. These alternative sentences were then spliced and edited as needed, and the authors take full responsibility for the content of the published article.

## Acknowledgments

This work was funded by the Fonds de recherche du Québec – Santé (FRQS) (269182). FRC and LDM are supported by the Fonds de recherche du Québec – Santé (FRQS) (B2X). KK was supported by the Natural Sciences and Engineering Research Council of Canada (NSERC) (USRA). We thank Rami Antoun and Madeleine Woisin for their help with Oct-TriA_1_ synthesis.

