## Supplementary Information for "*De novo* evolution of antibiotic resistance to Oct-TriA_1_"

Supplementary Material

**Authors**

Farhan R. Chowdhury*^a^, Laura Domínguez Mercado*^b^, Katya Kharitonov^a^, Brandon L. Findlay**^a,b^

^a^ Department of Biology, Concordia University, Montréal, Québec, Canada H4B 1R6

^b^ Department of Chemistry and Biochemistry, Concordia University, Montréal, Québec, Canada H4B 1R6

*These authors contributed equally to this work.

**This PDF file includes:**

Supplementary Tables 1-3

Supplementary Figures 1-5

**Supplementary Table 1. Mutations observed in SAGE isolates.**

| Strain | Position | Mutation | Annotation | Gene | Description |
| --- | --- | --- | --- | --- | --- |
| A26 | 1563034 | IS*2* (–) +5 bp | coding (43‑47/1320 nt) | *yddW* ← | liprotein, glycosyl hydrolase homolog |
| A26 | 3206481 | G→A | E26K (GAG→AAG) | *rpoD* → | RNA polymerase, sigma 70 (sigma D) factor |
| A26 | 3528471 | G→T | R253S (CGC→AGC) | *envZ* ← | sensory histidine kinase in two‑component regulatory system with OmpR |
| A51 | 55429 | C→T | G561S (GGT→AGT) | *lptD* ← | LPS assembly OM complex LptDE, beta‑barrel component |
| A51 | 56403 | (GTG)_3→2_ | coding (705‑707/2355 nt) | *lptD* ← | LPS assembly OM complex LptDE, beta‑barrel component |
| A51 | 572569 | (T)_6→7_ | intergenic (‑288/‑285) | *nmpC* ← / → *essD* | DLP12 prophage; truncated outer membrane porin (pseudogene);IS, phage, Tn; Phage or Prophage Related; outer membrane porin protein; locus of qsr prophage/DLP12 prophage; putative phage lysis protein |
| A51 | 1563034 | IS*2* (–) +5 bp | coding (43‑47/1320 nt) | *yddW* ← | liprotein, glycosyl hydrolase homolog |
| A51 | 1999224 | T→A | W11R (TGG→AGG) | *fliT* → | putative flagellar synthesis and assembly chaperone |
| A51 | 2306089 | T→C | N47S (AAC→AGC) | *ompC* ← | outer membrane porin protein C |
| A51 | 2399791 | IS*2* (–) +5 bp | coding (326‑330/939 nt) | *lrhA* ← | transcriptional repressor of flagellar, motility and chemotaxis genes |
| A51 | 3206481 | G→A | E26K (GAG→AAG) | *rpoD* → | RNA polymerase, sigma 70 (sigma D) factor |
| A51 | 3775555 | +A | coding (368/822 nt) | *cysE* ← | serine acetyltransferase |
| A51 | 3931636 | C→T | T17I (ACA→ATA) | *rbsR* → | transcriptional repressor of ribose metabolism |
| X34 | 55429 | C→T | G561S (GGT→AGT) | *lptD* ← | LPS assembly OM complex LptDE, beta‑barrel component |
| X34 | 240136 | T→C | L36P (CTG→CCG) | *gmhA* → | D‑sedoheptulose 7‑phosphate isomerase |
| X34 | 391960 | C→A | intergenic (‑217/‑135) | *ampH* ← / → *sbmA* | D‑alanyl‑D‑alanine‑ carboxypeptidase/endopeptidase; penicillin‑binding protein; weak beta‑lactamase/microcin B17 transporter |
| X34 | 1563034 | IS*2* (–) +5 bp | coding (43‑47/1320 nt) | *yddW* ← | liprotein, glycosyl hydrolase homolog |
| X34 | 1605138 | C→A | intergenic (‑201/+26) | *uxaB* ← / ← *yneF* | altronate oxidoreductase, NAD‑dependent/putative membrane‑bound diguanylate cyclase |
| X34 | 3206481 | G→A | E26K (GAG→AAG) | *rpoD* → | RNA polymerase, sigma 70 (sigma D) factor |
| X34 | 3512437 | IS*5* (+) +4 bp | intergenic (‑14/+384) | *aroK* ← / ← *hofQ* | shikimate kinase I/DNA catabolic putative fimbrial transporter |
| X34 | 4390177 | +T | coding (73/309 nt) | *hfq* → | global sRNA chaperone; HF‑I, host factor for RNA phage Q beta replication |
| XAM CM | 20771 | C→A | intergenic (‑263/+44) | *insA* ← / ← *rpsT* | IS1 repressor TnpA/30S ribosomal subunit protein S20 |
| XAM CM | 55428 | C→T | G561D (GGT→GAT) | *lptD* ← | LPS assembly OM complex LptDE, beta‑barrel component |
| XAM CM | 195916 | T→A | L501Q (CTG→CAG) | *bamA* → | BamABCDE complex OM biogenesis outer membrane pore‑forming assembly factor |
| XAM CM | 1563034 | IS*2* (–) +5 bp | coding (43‑47/1320 nt) | *yddW* ← | liprotein, glycosyl hydrolase homolog |
| XAM CM | 1972967 | IS*5* (+) +4 bp | intergenic (‑513/‑264) | *flhD* ← / → *uspC* | flagellar class II regulon transcriptional activator, with FlhC/universal stress protein |
| XAM CM | 2394241 | Δ7,687 bp |  | [*nuoF*]–*[alaA]* | [nuoF], *nuoE*, *nuoC*, *nuoB*, *nuoA*, *lrhA*, *[alaA]* |
| XAM CM | 2458311 | C→T | W59* (TGG→TAG) | *mlaA* ← | ABC transporter maintaining OM lipid asymmetry, OM lipoprotein component |
| XAM CM | 3206481 | G→A | E26K (GAG→AAG) | *rpoD* → | RNA polymerase, sigma 70 (sigma D) factor |
| XAM CM | 3674947 | +CA | coding (354/1497 nt) | *yhjJ* ← | putative periplasmic M16 family chaperone |
| XAM CM | 3797305 | IS*5* (–) +4 bp | coding (190‑193/1080 nt) | *waaB* ← | UDP‑D‑galactose:(glucosyl)lipopolysaccharide‑1, 6‑D‑galactosyltransferase |
| XAM_WT | 65,196 | +C | *coding (585/2352 nt)* | *polB* ← | DNA polymerase II |
| XAM_WT | 1,652,589 | C→A | *Y88* (TAC→TAA)* | *ynfE* → | putative selenate reductase, periplasmic |
| XAM_WT | 1,972,715 | G→T | *intergenic (‑261/‑519)* | *flhD* ← / → *uspC* | flagellar class II regulon transcriptional activator, with FlhC/universal stress protein |
| XAM_WT | 4,171,661 | A→T | *K163N (AAA→AAT)* | *rpoB* → | RNA polymerase, beta subunit |

**Table S2: Gene ontology groups from all of the identified mutations of the evolved strains determined with ShinyGo v0.741(69).**

| **Enrichment FDR** | **N^1^** | **Pathway Genes** | **Fold Enrichment** | **Pathway** | **URL** | **Genes** |
| --- | --- | --- | --- | --- | --- | --- |
| **1.95156790441107E-05** | 5 | 16 | 41.6028225806452 | Respiratory chain complex i | <http://amigo.geneontology.org/amigo/term/GO:0045271> | nuoF nuoE nuoC nuoB nuoA |
| **1.95156790441107E-05** | 5 | 16 | 41.6028225806452 | Plasma membrane respiratory chain complex i | <http://amigo.geneontology.org/amigo/term/GO:0045272> | nuoF nuoE nuoC nuoB nuoA |
| **1.95156790441107E-05** | 5 | 14 | 47.5460829493088 | Quinone |  | nuoF nuoE nuoC nuoB nuoA |
| **2.06263497889355E-05** | 5 | 17 | 39.1555977229602 | NADH dehydrogenase complex | <http://amigo.geneontology.org/amigo/term/GO:0030964> | nuoF nuoE nuoC nuoB nuoA |
| **9.70169447122268E-05** | 5 | 25 | 26.6258064516129 | Quinone binding | <http://amigo.geneontology.org/amigo/term/GO:0048038> | nuoF nuoE nuoC nuoB nuoA |
| **9.70169447122268E-05** | 5 | 24 | 27.7352150537634 | Plasma membrane respirasome | <http://amigo.geneontology.org/amigo/term/GO:0070470> | nuoF nuoE nuoC nuoB nuoA |
| **9.70169447122268E-05** | 5 | 25 | 26.6258064516129 | Respiratory chain complex | <http://amigo.geneontology.org/amigo/term/GO:0098803> | nuoF nuoE nuoC nuoB nuoA |
| **0.000114385296756409** | 4 | 12 | 44.3763440860215 | Quinone |  | nuoF nuoE nuoB nuoA |
| **0.000114385296756409** | 5 | 27 | 24.6535244922342 | Respirasome | <http://amigo.geneontology.org/amigo/term/GO:0070469> | nuoF nuoE nuoC nuoB nuoA |
| **0.000147921974039749** | 4 | 13 | 40.9627791563275 | Ubiquinone |  | nuoF nuoE nuoC nuoA |
| **0.000438316641715869** | 4 | 17 | 31.3244781783681 | NADH dehydrogenase (ubiquinone) activity | <http://amigo.geneontology.org/amigo/term/GO:0008137> | nuoF nuoC nuoB nuoA |
| **0.000513881818946486** | 4 | 18 | 29.584229390681 | NADH dehydrogenase (quinone) activity | <http://amigo.geneontology.org/amigo/term/GO:0050136> | nuoF nuoC nuoB nuoA |
| **0.000743211509090046** | 4 | 20 | 26.6258064516129 | Ubiquinone, and fumarate reductase complex |  | nuoF nuoE nuoB nuoA |
| **0.000848042580787152** | 4 | 21 | 25.357910906298 | NADH dehydrogenase activity | <http://amigo.geneontology.org/amigo/term/GO:0003954> | nuoF nuoC nuoB nuoA |
| **0.00163817213050132** | 4 | 25 | 21.3006451612903 | Oxidative phosphorylation |  | nuoF nuoE nuoB nuoA |
| **0.00244694298116853** | 4 | 28 | 19.0184331797235 | Oxidoreductase activity, acting on nad(p)h, quinone or similar compound as acceptor | <http://amigo.geneontology.org/amigo/term/GO:0016655> | nuoF nuoC nuoB nuoA |
| **0.00400197596550932** | 5 | 62 | 10.7362122788762 | Oxidoreductase complex | <http://amigo.geneontology.org/amigo/term/GO:1990204> | nuoF nuoE nuoC nuoB nuoA |
| **0.00639200607085923** | 7 | 160 | 5.82439516129032 | Membrane protein complex | <http://amigo.geneontology.org/amigo/term/GO:0098796> | bamA ompC nuoF nuoE nuoC nuoB nuoA |
| **0.00734355120593817** | 5 | 72 | 9.24507168458781 | Translocase |  | nuoF nuoE nuoC nuoB nuoA |
| **0.0112586016181981** | 6 | 125 | 6.3901935483871 | Cell outer membrane | <http://amigo.geneontology.org/amigo/term/GO:0009279> | lptD bamA yddW ompC mlaA hofQ |
| **0.011365060447705** | 4 | 44 | 12.1026392961877 | Respirasome, and nitrate assimilation |  | nuoF nuoE nuoB nuoA |
| **0.0148326303699135** | 6 | 135 | 5.9168458781362 | Outer membrane | <http://amigo.geneontology.org/amigo/term/GO:0019867> | lptD bamA yddW ompC mlaA hofQ |
| **0.0148326303699135** | 6 | 136 | 5.87333965844402 | External encapsulating structure | <http://amigo.geneontology.org/amigo/term/GO:0030312> | lptD bamA yddW ompC mlaA hofQ |
| **0.0148326303699135** | 6 | 135 | 5.9168458781362 | Macromolecule localization | <http://amigo.geneontology.org/amigo/term/GO:0033036> | lptD bamA sbmA ompC mlaA hofQ |
| **0.0169037765170519** | 4 | 52 | 10.2406947890819 | Oxidoreductase activity, acting on nad(p)h | <http://amigo.geneontology.org/amigo/term/GO:0016651> | nuoF nuoC nuoB nuoA |
| **0.0169037765170519** | 6 | 142 | 5.62517037710132 | NAD |  | uxaB nuoF nuoE nuoC nuoB nuoA |
| **0.0169037765170519** | 5 | 92 | 7.23527349228612 | Cell outer membrane |  | lptD bamA ompC mlaA hofQ |
| **0.0192931642820252** | 6 | 147 | 5.43383805134957 | Catalytic complex | <http://amigo.geneontology.org/amigo/term/GO:1902494> | nuoF nuoE nuoC nuoB nuoA cysE |
| **0.0215782433841346** | 2 | 6 | 44.3763440860215 | Phospholipid transport | <http://amigo.geneontology.org/amigo/term/GO:0015914> | ompC mlaA |
| **0.0215782433841346** | 2 | 6 | 44.3763440860215 | Regulation of organelle assembly | <http://amigo.geneontology.org/amigo/term/GO:1902115> | flhD fliT |

^1^ N = number of genes.

**Table S3: Gene ontology groups, high level GO categories, from all of the identified mutations of the evolved strains determined with ShinyGo v0.741(69).**

| N^1^ | High level GO category | Genes |
| --- | --- | --- |
| 27 | Cellular process | *insA1 rpsT lptD bamA gmhA ampH sbmA essD yddW uxaB yneF flhD uspC fliT ompC nuoF nuoE nuoB lrhA alaA mlaA rpoD aroK hofQ envZ cysE waaB* |
| 21 | Binding | *rpsT gmhA ampH sbmA yneF flhD ompC nuoF nuoE nuoC nuoB nuoA lrhA alaA rpoD aroK hofQ envZ yhjJ rbsR hfq* |
| 21 | Metabolic process | *insA1 rpsT gmhA ampH yddW uxaB flhD nuoF nuoE nuoC nuoB nuoA lrhA alaA rpoD aroK hofQ envZ yhjJ cysE waaB* |
| 16 | Catalytic activity | *gmhA ampH yddW uxaB yneF nuoF nuoE nuoC nuoB nuoA alaA aroK envZ yhjJ cysE waaB* |
| 16 | Cellular metabolic process | *insA1 rpsT gmhA uxaB flhD nuoF nuoE nuoB lrhA alaA rpoD aroK hofQ envZ cysE waaB* |
| 15 | Membrane | *lptD bamA ampH sbmA yddW yneF ompC nuoF nuoE nuoC nuoB nuoA mlaA hofQ envZ* |
| 15 | Primary metabolic process | *insA1 rpsT gmhA ampH uxaB flhD lrhA alaA rpoD aroK hofQ envZ yhjJ cysE waaB* |
| 15 | Organic substance metabolic process | *insA1 rpsT gmhA ampH uxaB flhD lrhA alaA rpoD aroK hofQ envZ yhjJ cysE waaB* |
| 15 | Cell periphery | *lptD bamA ampH sbmA yddW yneF ompC nuoF nuoE nuoC nuoB nuoA mlaA hofQ envZ* |
| 15 | Organic cyclic compound binding | *rpsT ampH sbmA yneF flhD nuoF nuoC lrhA alaA rpoD aroK hofQ envZ rbsR hfq* |
| 15 | Heterocyclic compound binding | *rpsT ampH sbmA yneF flhD nuoF nuoC lrhA alaA rpoD aroK hofQ envZ rbsR hfq* |
| 13 | Intracellular | *rpsT gmhA uxaB flhD uspC fliT nuoC alaA rpoD aroK cysE waaB hfq* |
| 12 | Nitrogen compound metabolic process | *insA1 rpsT ampH flhD lrhA alaA rpoD aroK hofQ envZ yhjJ cysE* |
| 12 | Ion binding | *gmhA ampH sbmA yneF ompC nuoF nuoE nuoB alaA aroK envZ yhjJ* |
| 10 | Protein-containing complex | *rpsT lptD bamA ompC nuoF nuoE nuoC nuoB nuoA cysE* |
| 10 | Biological regulation | *rpsT ampH yneF flhD fliT lrhA rpoD envZ rbsR hfq* |
| 9 | Regulation of biological process | *ampH yneF flhD fliT lrhA rpoD envZ rbsR hfq* |
| 9 | Response to stimulus | *lptD sbmA uspC ompC alaA rpoD aroK envZ cysE* |
| 9 | Cellular component organization or biogenesis | *rpsT lptD bamA gmhA ampH yddW flhD fliT mlaA* |
| 9 | Plasma membrane | *ampH sbmA yneF nuoF nuoE nuoC nuoB nuoA envZ* |
| 9 | Biosynthetic process | *rpsT gmhA flhD lrhA alaA rpoD aroK cysE waaB* |
| 9 | Cellular component organization | *rpsT lptD bamA gmhA ampH yddW flhD fliT mlaA* |
| 8 | Localization | *lptD bamA sbmA fliT ompC nuoB mlaA hofQ* |
| 8 | Small molecule binding | *ampH sbmA yneF nuoF nuoC alaA aroK envZ* |
| 8 | Regulation of cellular process | *yneF flhD fliT lrhA rpoD envZ rbsR hfq* |
| 7 | Establishment of localization | *lptD bamA sbmA ompC nuoB mlaA hofQ* |
| 7 | Membrane protein complex | *bamA ompC nuoF nuoE nuoC nuoB nuoA* |
| 6 | Oxidoreductase activity | *uxaB nuoF nuoE nuoC nuoB nuoA* |
| 6 | Transferase activity | *yneF alaA aroK envZ cysE waaB* |
| 6 | Outer membrane | *lptD bamA yddW ompC mlaA hofQ* |
| 6 | External encapsulating structure | *lptD bamA yddW ompC mlaA hofQ* |
| 6 | Intrinsic component of membrane | *bamA sbmA yneF ompC nuoA envZ* |
| 6 | Envelope | *lptD bamA yddW ompC mlaA hofQ* |
| 6 | Macromolecule localization | *lptD bamA sbmA ompC mlaA hofQ* |
| 6 | Carbohydrate derivative binding | *gmhA sbmA yneF nuoF aroK envZ* |
| 5 | Regulation of metabolic process | *flhD lrhA rpoD rbsR hfq* |
| 5 | Cellular component biogenesis | *rpsT lptD bamA gmhA flhD* |
| 5 | Respirasome | *nuoF nuoE nuoC nuoB nuoA* |
| 5 | Oxidoreductase complex | *nuoF nuoE nuoC nuoB nuoA* |
| 4 | Molecular function regulator | *rpsT lrhA rpoD rbsR* |
| 4 | Response to stress | *uspC ompC alaA rpoD* |
| 4 | Hydrolase activity | *ampH yddW envZ yhjJ* |
| 4 | Response to chemical | *lptD sbmA alaA aroK* |
| 4 | Positive regulation of biological process | *flhD lrhA rbsR hfq* |
| 4 | Negative regulation of biological process | *yneF fliT lrhA hfq* |
| 4 | Cellular response to stimulus | *uspC ompC alaA envZ* |
| 4 | Regulation of molecular function | *rpsT lrhA rpoD rbsR* |
| 3 | Biological adhesion | *bamA yneF ompC* |
| 3 | DNA-binding transcription factor activity | *lrhA rpoD rbsR* |
| 3 | Catabolic process | *ampH uxaB hofQ* |
| 3 | Response to abiotic stimulus | *uspC rpoD cysE* |
| 3 | Metal cluster binding | *nuoF nuoE nuoB* |
| 2 | Transporter activity | *sbmA ompC* |
| 2 | Protein binding | *gmhA envZ* |
| 2 | Cell adhesion | *bamA yneF* |
| 2 | Transmembrane transporter activity | *sbmA ompC* |
| 2 | Regulation of cellular component biogenesis | *flhD fliT* |
| 2 | Interspecies interaction between organisms | *essD ompC* |
| 2 | Regulation of biological quality | *ampH hfq* |
| 2 | Cell wall organization or biogenesis | *ampH yddW* |
| 2 | Side of membrane | *bamA envZ* |
| 1 | Cell killing | *essD* |
| 1 | Structural molecule activity | *rpsT* |
| 1 | Signaling | *envZ* |
| 1 | Locomotion | *fliT* |
| 1 | Organelle | *rpsT* |
| 1 | Cell aggregation | *yneF* |
| 1 | Structural constituent of ribosome | *rpsT* |
| 1 | Protein folding | *fliT* |
| 1 | Drug binding | *ampH* |
| 1 | Response to external stimulus | *uspC* |
| 1 | Carbon utilization | *hofQ* |
| 1 | Isomerase activity | *gmhA* |
| 1 | Toxin transmembrane transporter activity | *sbmA* |
| 1 | Cytolysis | *essD* |
| 1 | Enzyme regulator activity | *rpsT* |
| 1 | Killing of cells of other organism | *essD* |
| 1 | Regulation of localization | *yneF* |
| 1 | Amide binding | *ampH* |
| 1 | Regulation of locomotion | *yneF* |
| 1 | Periplasmic space | *yhjJ* |
| 1 | Xenobiotic transmembrane transporter activity | *sbmA* |
| 1 | Non-membrane-bounded organelle | *rpsT* |
| 1 | Intracellular organelle | *rpsT* |
| 1 | Adhesion of symbiont to host | *ompC* |
| 1 | Regulation of response to stimulus | *envZ* |
| 1 | Cell motility | *fliT* |
| 1 | Regulation of developmental process | *ampH* |
| 1 | Cellular localization | *bamA* |
| 1 | Localization of cell | *fliT* |
| 1 | Intraspecies interaction between organisms | *yneF* |
| 1 | Molecular adaptor activity | *uspC* |
| 1 | Aggregation of unicellular organisms | *yneF* |
| 1 | Sulfur compound binding | *ampH* |
| 1 | Ribonucleoprotein complex | *rpsT* |

^1^ N = number of genes.

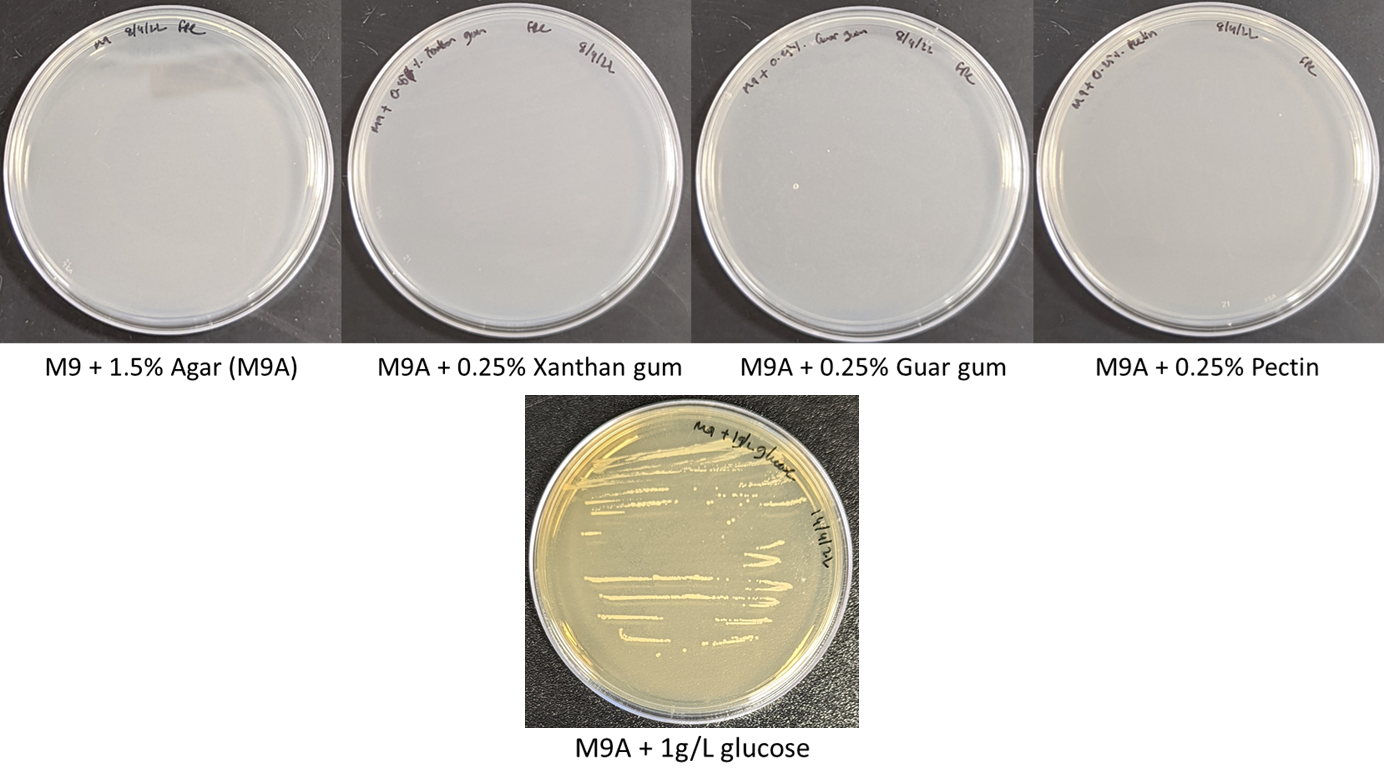

**Supplementary Figure 1. Growth on gelling agents.** No growth was observed when *E. coli* cells were streaked on plates made with M9 + 1.5% agar (M9A) or M9A supplemented with xanthan gum, guar gum, or pectin at 0.25% w/v following 24 h of incubation. M9A supplemented with 1 g/L of glucose showed clear growth.

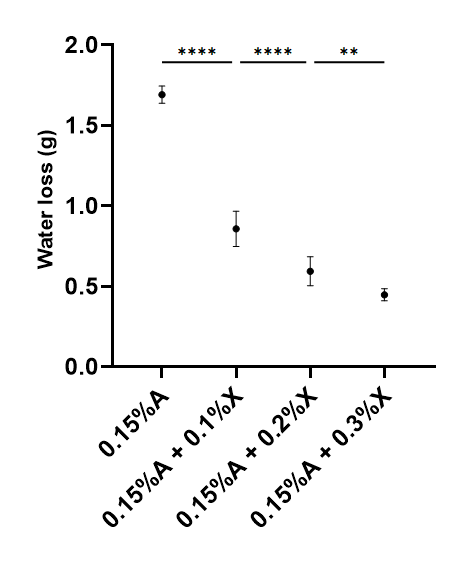

**Supplementary Figure 2. Synaeresis at various xanthan gum concentrations.** Increasing the amount of xanthan gum added to 0.15% agar medium reduces water loss, with diminishing returns at higher xanthan gum concentrations. A = agar. X = Xanthan gum. *p < 0.05, **p < 0,01, *** p < 0.001, ****p < 0.0001, one-way ANOVA with Fisher’s LSD test.

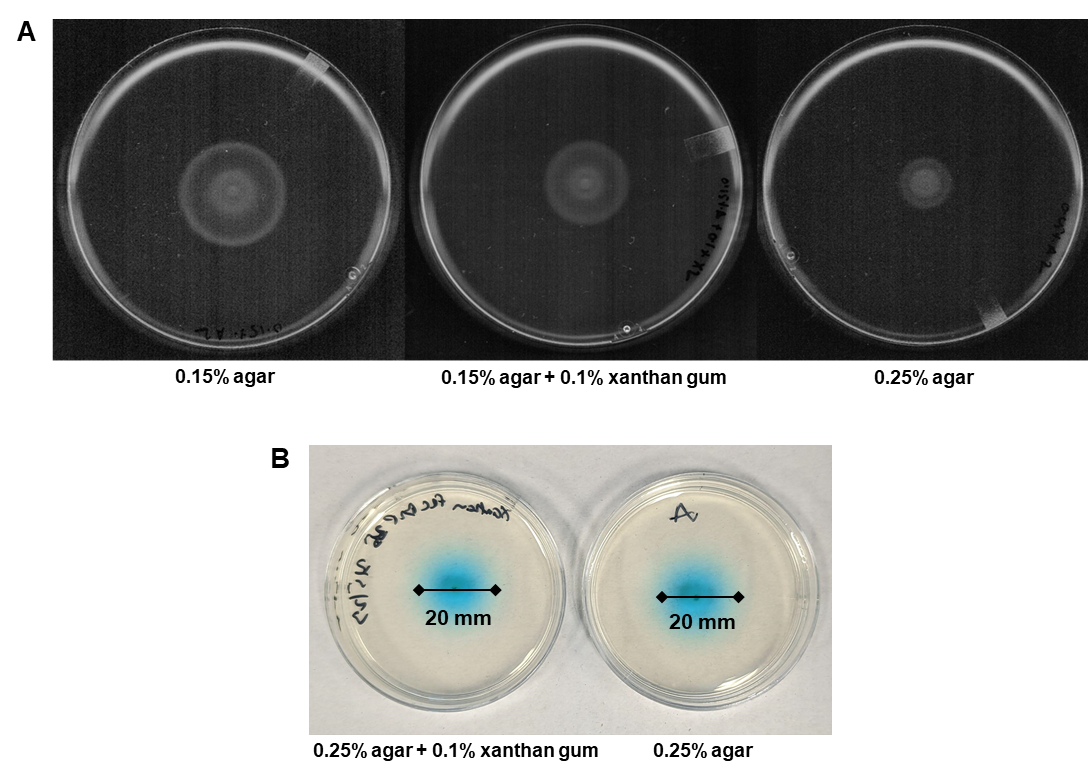

**Supplementary Figure 3. Motility and diffusion assays.** (A) Cells swimming through 0.15% agar laced with 0.1% xanthan gum show increased motility relative to 0.25 % agar. Representative pictures from three independent replicates are shown. (B) Diffusion rate of malachite green in 0.25% agar and 0.15% agar + 0.1% xanthan gum are similar. 2 μL of malachite green was impregnated in the center of each plate (60 x 15 mm) made with 10 mL of media. Measurements were taken after 8 h.

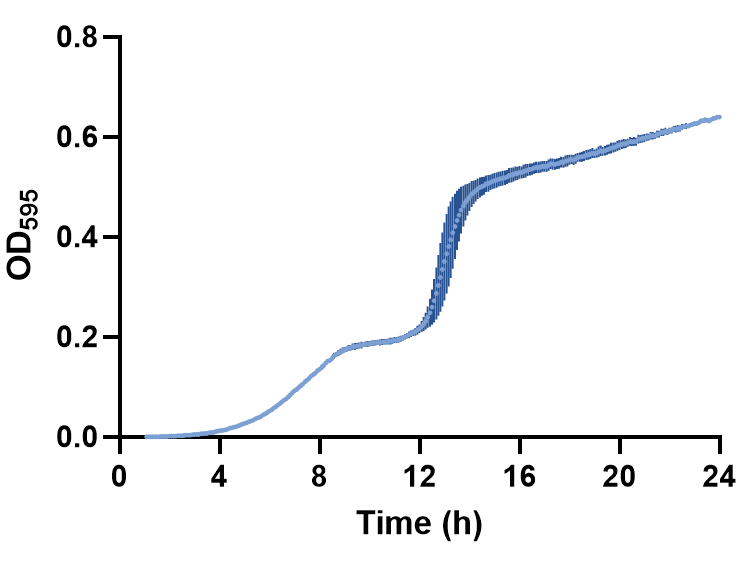

**Supplementary Figure 4. Growth of XAM CM in MHB.** Cells show a diauxic growth curve. The mean of three independent replicates is shown. Error bars indicate standard deviation.

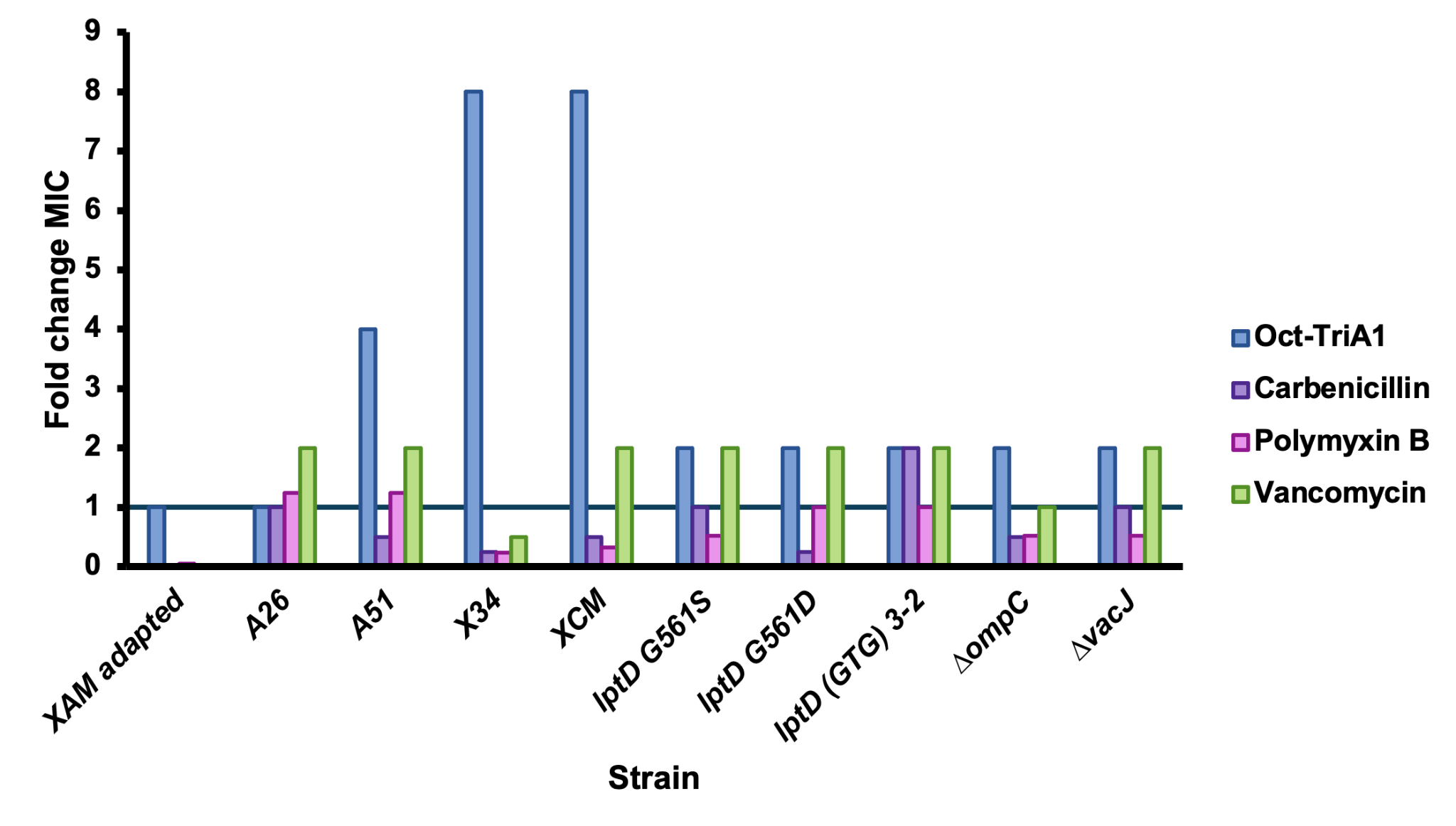

**Supplementary Figure 5. Fold change MIC of the evolved and mutated strains.** Fold change MIC values for Oct-TriA1 and other membrane targeting antibiotics compared to the ancestral strain.
